# Visualizing histone H4K20me1 in knock-in mice expressing the mCherry-tagged modification-specific intracellular antibody

**DOI:** 10.1101/2024.03.13.584715

**Authors:** Yuko Sato, Maoko Takenoshita, Miku Ueoka, Jun Ueda, Kazuo Yamagata, Hiroshi Kimura

## Abstract

During development and differentiation, histone modifications dynamically change locally and globally, associated with transcriptional regulation, DNA replication and repair, and chromosome condensation. The level of histone H4 Lys20 monomethylation (H4K20me1) increases during the G2 to M phases of the cell cycle and is enriched in facultative heterochromatin, such as inactive X chromosomes in cycling cells. To track the dynamic changes of H4K20me1 in living cells, we have developed a genetically encoded modification-specific intracellular antibody (mintbody) probe that specifically binds to the modification. Here, we report the generation of knock-in mice in which the coding sequence of the mCherry-tagged version of the H4K20me1-mintbody is inserted into the *Rosa26* locus. The knock-in mice, which ubiquitously expressed the H4K20me1-mintbody, developed normally and were fertile, indicating that the expression of the probe does not disturb the cell growth, development, or differentiation. Various tissues isolated from the knock-in mice exhibited nuclear fluorescence without the need for fixation. The H4K20me1-mintbody was enriched in inactive X-chromosomes in developing embryos and in XY bodies during spermatogenesis. The knock-in mice will be useful for the histochemical analysis of H4K20me1 in any cell types.

## Introduction

Posttranslational histone modifications play critical roles in development and differentiation through the regulation of genome functions including gene expression, DNA replication, and genome maintenance (Millán-Zambrano et al. 2022; Weinzapfel et al. 2024). Some modifications, such as histone H3 Lys9 trimethylation (H3K9me3) and Lys27 trimethylation (H3K27me3), are maintained through cell generations to epigenetically sustain a transcriptionally repressive chromatin state (Margueron and Reinberg 2010). Other modifications exhibit dynamic changes upon gene activation, like H3 Lys27 acetylation (H3K27ac) (Katan-Khaykovich and Struhl 2002; Stasevich et al. 2014), or fluctuate during the cell cycle, like H3 Ser10 phosphorylation (H3S10ph), which occurs on mitotic chromosomes (Hendzel et al. 1997; Johansen and Johansen 2006; Hayashi-Takanaka et al. 2020). The level of H4 Lys20 monomethylation (H4K20me1) also changes drastically during the cell cycle, with an increase during the G2 phase and a decrease during the subsequent G1 phase (Rice et al. 2002; Pesavento et al. 2008; van Nuland and Gozani 2015; Sato et al. 2016). H4K20me1 is generally distributed across the gene body and on inactive X chromosomes, which are typical of facultative heterochromatin, in cycling cells (Kohlmaier et al. 2004; Calabrese et al. 20212; Tjalsma et al. 2021). In addition, H4K20me1 is involved in DNA damage repair (Schotta et al 2008; Oda et al., 2009; Dulev 2014; Jørgensen et al. 2013) and centromere chromatin formation (Hori et al. 2014). However, the precise function and regulation of H4K20me1 have yet to be fully understood (Corvalan AZ, Coller HA 2021).

To track the changes in the levels and intranuclear localization of histone modifications during the cell cycle and during differentiation, we have developed a genetically encoded modification-specific intracellular antibody (mintbody) probe, which is the single-chain variable fragment (scFv) of a specific antibody tagged with a fluorescent protein (Kimura et al. 2015; Sato et al. 2021). Using the H4K20me1-specific mintbody (H4K20me1-mintbody), we have demonstrated H4K20me1’s cell cycle oscillation in HeLa cells and accumulation on inactivating X chromosomes in differentiating embryonic stem cells (ESCs) (Sato et al. 2016; Tjalsma et al. 2021). The mintbody probes can be expressed in model organisms, including yeast, nematodes, flies, frogs, and plants, indicating that mintbody expression does not affect development, differentiation, and fertility (Sato et al. 2021). However, the expression of mintbody in mammals has never been demonstrated. Here, we generated mice in which the H4K20me1-mintbody (a red fluorescent protein, mCherry version) is knocked into the *Rosa26* locus (Soriano et al. 1999). Homozygous knock-in mice that exhibit H4K20me1-mintbody expression in various tissues developed normally and were fertile. The mintbody-expressing mice will be useful for visualizing and tracking the specific modification in any given cell types in live.

## Materials and Methods

### Generation of mouse embryonic stem cells and knock-in mice

Mouse care and experimental procedures were approved by the Institutional Animal Experiment Committee of the Tokyo Institute of Technology and the Animal Care and Use Committee of the Research Institute for Microbial Diseases, Osaka University. All animal experiments were conducted in accordance with institutional and governmental guidelines. Mice were maintained on a 12:12 dark: light cycle at a constant temperature of 22–23°C in ventilated cages and were housed in a specific pathogen-free facility with free access to food and water.

Knock-in mouse C57BL/6N ESCs were generated using H4K20me1-mintbody (clone 15F11, mCherry version; Sato et al. 2016), as described previously (Sato et al. 2013; Ueda et al. 2017). Briefly, The FRT-Neo^r^-H4K20me1-mintbody (mCherry)-FRT cassette, subcloned in the pBigT vector (Srinivas et al. 2001), was inserted into the ROSA26 vector (Soriano et al. 1999). ESCs were electroporated with the linearized vector using XhoI and selected in 150 μg/mL G418 (Thermo Fisher Scientific). Clones were validated by genomic PCR using the following primers; ROSA26-SA-Fw, 5′-CCTAAAGAAGAGGCTGTGCTTTGG-3′; ROSA26-LA-Rev, 5′-GTAGTTACTCCACTTTCAAGTTCCTTATAA-3′; 15F11-scFv-183as, 5′-AGTGTTTGGATAGTAGGTATAACTACCAC-3′; and mCherry-C1-F-1245S 5′-GGACTACACCATCGTGGAAC-3′. The resulting 15F11-mintbody knock-in ESCs were injected into blastocysts, which were then transferred to the uterus of day 2.5 pseudopregnant mothers to generate chimeric mice. Knock-in alleles in mice were assessed by genomic PCR using ROSA26-NotI-Fw, 5’-GAGCGGCCGCCCACCCTCCCCTTCCTCTGG-3' and ROSA26-NruI-Rev, 5’-CCTCGCGACACTGTATTTCATACTGTAGTA-3’ primers. Sex determination was performed by genomic PCR using Y-specific SRY primers (SRY-F, 5’-CTGTGTAGGATCTTCAATCTCT-3’; and SRY-R, 5’-GTGGTGAGAGGCACAAGTTGGC-3’) and Ube1X primers(Ube IX-F, 5’-TGGTCTGGACCCAAACGCTGTCCACA-3’; and Ube1X-R, 5’-GGCAGCAGCCATCACATAATCCAGATG-3’) to yield PCR products with different sized bands for X (217 bp) and Y (198 bp) chromosomes (Chuma and Nakatsuji 2001).

### Live-cell microscopy

Mouse tissue and embryos expressing H4K20me1-mintbody were placed onto a 35-mm glass bottom dish (No. 1.5, MatTek) and imaged using a confocal microscope, either a point scan system (Olympus FV-1000 with IX-83, or Nikon A1 with Ti-2), or a spinning disk system (Yokogawa CSU-W1 with Nikon Ti-E). Mouse tissues, dissected using a knife, and preimplantation embryos were imaged using an Olympus FV-1000, operated by the built-in FV1000 software (ver. 3.1a), with a 30× UPlanSApo silicone-immersion objective lens (NA 1.05), a 405/488/543/633 dichroic mirror, and a BA575-675 bandpass filter, using the following settings: 800 × 800 pixels; pinhole 75 μm; 3.0× zoom; 4.0 μs/pixel; 4× averaging; 22% transmission of a 2 mW HeNe 543-nm laser. Mouse E6.5 and E7.5 embryos were imaged using a Yokogawa CSU-W1, operated by NIS-Elements AR (ver. 4.3; Nikon), with a 40× CFI Apo Lambda S water-immersion objective lens (NA 1.25), a 405/488/561/640 dichroic mirror, a 590LP emission filter, a laser unit (Nikon LU-N4; 100% transmission of the 561-nm laser line), and EM-CCD (Andor iXon3; conversion gain 5x; gain multiplier 300; exposure time 500 ms). Testis samples were imaged using a Nikon A1, operated by NIS-Elements AR (ver. 5.21; Nikon), with a 25× Plan Apo Lambda S silicone-immersion objective lens (NA 1.05), a 405/488/561/640 dichroic mirror, and a 595/50 emission filter, and a laser unit (Nikon LU-N4; 1% transmission of the 561-nm laser line), using the following settings: 1024 × 1024 pixels; pinhole 21.7 μm; 0.72× zoom; 32 μs/pixel; 8× averaging). Seminiferous tubules were imaged using a Yokogawa CSU-W1, operated by NIS-Elements AR (ver. 4.3; Nikon) as above, except using an Olympus 30× UPlanSApo silicone-immersion objective lens (NA 1.05),

### Immunofluorescence

All procedures were performed at room temperature, unless stated otherwise. E7.5 embryos were fixed with 4% paraformaldehyde in phosphate-buffered saline (PBS; Wako) for 10 min, washed with PBS three times for 5 min each, and permeabilzed with 0.5% Trion X-100 (Fujifilm) and 0.5% Blocking One-P (nacalai tesque) in PBS for 10 min. After washing with PBS, embryos were incubated in 10 μg/mL Alexa Fluor 488-conjugated mouse anti-H3K27me3 (clone CMA323/1E7; Hayashi-Takanaka et al., 2011), 4 μg/mL Cy3-conjugated mouse anti-H4K20me1 (clone CMA421/15F11; Hayashi-Takanaka et al., 2015), and 0.1 μg/mL Hoechst 33342 (nacalai tesque) in PBS containing 10% Blocking One-P and 0.5% Tween 20 (Wako) for 1-3 days at 4°C. After washing with PBS three times for 3 min each, embryos were mounted on a 35-mm glass-bottom dish using 100-200 μL of 0.5% low-gelling temperature agarose (Sigma) in PBS prewarmed to 40°C. Before the agarose gel hardened, the position of the embryos was adjusted for imaging using a needle. Mouse tissue sections were prepared as previously described (Goto et al., 2023). The embryos were imaged using a Yokogawa CSU-W1 as described above using a 447/60, a 520/35, a 590LP emission filters, and 405-, 488-, and 561-nm laser lines (Nikon LU-4N).

Cells in seminiferous tubules were suspended in Dulbecco’s Modified Eagle’s Medium (DMEM), High Glucose (nacalai tesque), 10% Fetal Bovine Serum (Gibco; Thermo Fisher Scientific), 1% glutamine-penicillin-streptomycin solution (SIGMA). Approximately ∼2×10^5^ cells in 100 μL were spun (1,000 rpm; 2 min) onto a coverslip using a cytospin (Cytopro). Cells were fixed in 4% paraformaldehyde in 250 mM HEPES (pH7.4) containing 0.1% Triton X-100 for 5 min, washed with PBS three times, and permeabilized with 1% Triton X-100 in PBS for 20 min. After blocking with Blocking One-P for 20 min, cells were washed with PBS and incubated with the primary antibodies (0.1 μg/mL rabbit anti-SCP3, ab15093, abcam; and 2 μg/mL mouse anti-γH2AX, clone 20A8; Yamagata et al., 2019) in 10% Blocking One-P in PBS for 2 h. After washing with PBS three times for 5 min each, cells were incubated in the secondary antibodies (1 μg/mL Cy5-conjugated Donkey anti-rabbit IgG H+L, 711-175-152, Jackson ImmunoResearch; and 1 μg/mL Alexa Fluor 488-conjugated version of Donkey anti-mouse IgG H+L, 715-005-150, Jackson ImmunoResearch) and 0.1 μg/mL Hoechst 33342 in 10% Blocking One-P in PBS for 2 h. After washing with PBS three times for 5 min each, a coverslip was mounted to a glass slide using Prolong Diamond (Thermo Fisher Scientific). Confocal images were acquired using an Andor DragonFly with Nikon Ti-E, operated by Fusion 2.2.0.50 (Andor), with a 100× PlanApo oil-immersion objective lens (NA 1.4), a 405/488/561/640 dichroic mirror, a 445/46, a 521/38, an a 600/50 emission filters, a laser unit (Andor; LC-ILE-400-M; 10% transmission of 405-nm, 20% transmission of 488-nm, and 50% transmission of 561-nm), and EM-CCD (Andor; iXonUltra; gain multiplier 300; exposure time 500, 100, and 1,500 ms for 405-, 488-, 561-nm laser lines, respectively).

## Results and Discussion

### Generation of mice expressing H4K20me1-mintbody

To generate mice expressing the H4K20me1-mintbody, we established ESC lines in which a FLP recombinase-exercisable Neo-resistant (Neo^r^) gene cassette, flanking the H4K20me1-mintbody (mCherry version) coding sequence, was knocked into a *Rosa26* allele (Fig. 1a). From chimeric mice generated by microinjecting knock-in ESCs into blastocysts, two independent germline-transmitted lines were cloned and then crossed with mice expressing FLP recombinase to delete the Neo^r^ gene. The resulting homozygous H4K20me1-mintbody knock-in C57BL/6N mice (Fig. 1b) have been maintained over five years (tens of generations), indicating that H4K20me1-mintbody mice develop normally and were fertile.

**Fig. 1.**
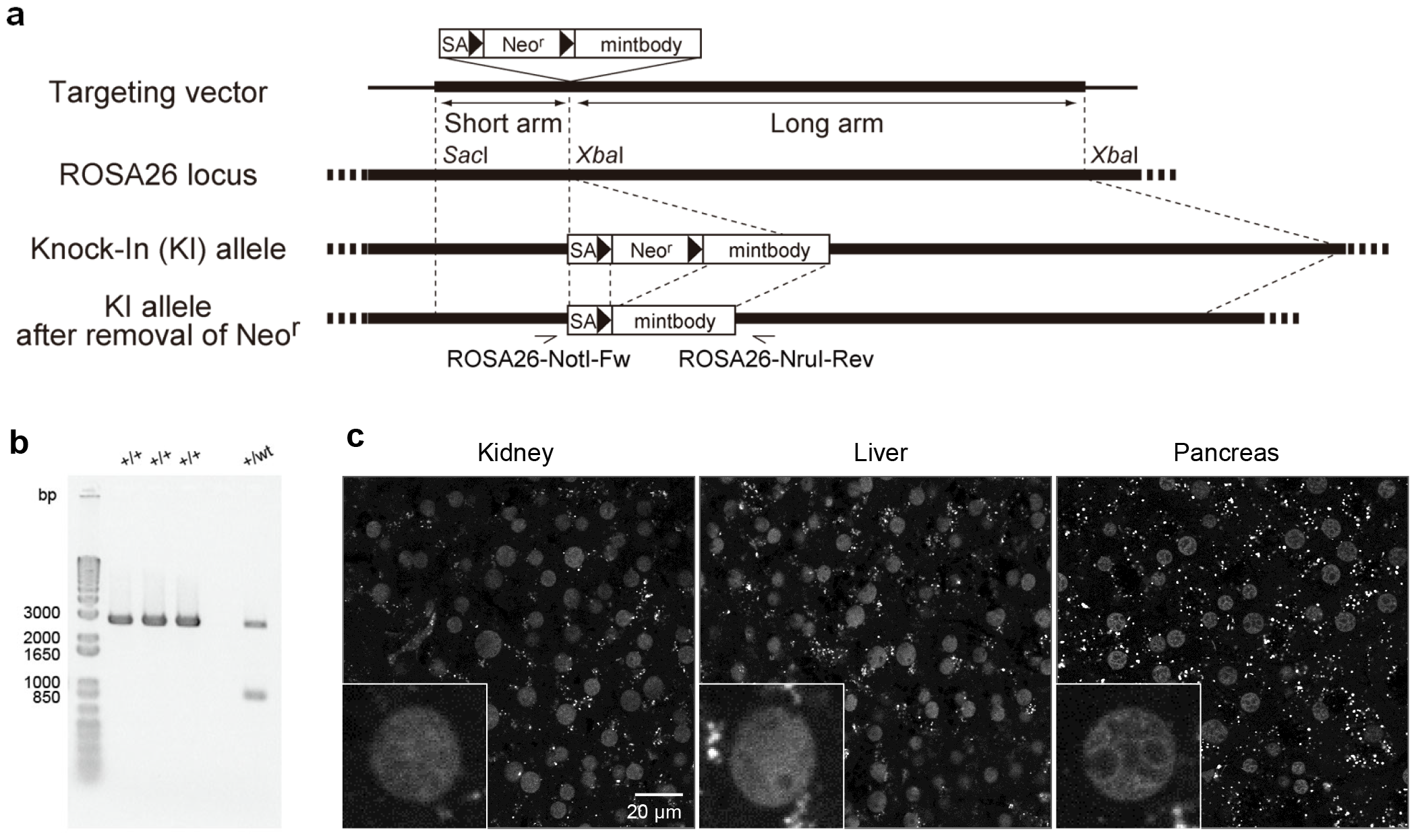
Generation of H4K20me1-mintbody knock-in mice. (a) Schematic drawing of the knock-in strategy. The targeting vector contains a splicing acceptor (SA), Neo resistant (Neo^r^) gene flanked by two FRT sites for excision by FLP recombinase, and the H4K20me1-mintbody positioned between the short and long arms of Rosa26 locus. Mouse ESCs harboring the knock-in allele were selected for Neo resistance. By crossing knock-in with FLP-expressing mice, H4K20me1-mintbody expression is achieved following the removal of Neo^r^. Sites for primers used to validate the knock-in and Neo^r^ removal are indicated. (b) Validation of homozygous knock-in using genomic PCR. Three homozygous and one heterozygous mice were analyzed. (c) H4K20me1-mintbody in various tissues (kidney, liver, and pancreas) of homozygous knock-in mice. Insets show magnified views of single nuclei.

We investigated whether H4K20me1-mintbody signals could be detected in knock-in mice. In a variety of tissues, H4K20me1-mintbody signals were detected without the need of fixation and thin-sectioning (Fig. 1c). Note that the distribution of H4K20me1-mintbody was the same in the two independent knock-in lines. In most cells, H4K20me1-mintbody exhibited little specific concentrations, consistent with the observation that H4K20me1 is not concentrated in constitutive heterochromatin. However, although H4K20me1 is known to be concentrated in the inactive X chromosome (Xi) in cultured cells, including differentiating ESCs, no Xi-like foci were observed in adult female mouse tissues (Fig. 1c). This suggests that H4K20me1 is no longer concentrated on Xi in non-dividing, differentiated cells, probably because monomethylation is converted to demethylation by Suv420H1 during the cell cycle arrest (Schotta et al 2008; van Nuland and Gozani 2015; Corvalan AZ, Coller HA 2021).

### H4K20me1-mintbody highlights inactive X chromosomes in embryos

To visualize the distribution of H4K20me1-mintbody during development, we prepared preimplantation and post-implantation embryos, from 3.5 days post coitum (dpc) blastocysts to E14.5. H4K20me1-mintbody was concentrated in single foci in nuclei in some blastocysts, while it was distributed more homogenously in others (Fig. 2a). As H4K20me1 is enriched in Xi in differentiating cells, this suggests that embryos with and without H4K20me1-mintbody foci likely represent female and male, respectively. In some of E6.5 and E7.5 post-implantation embryos, most nuclei exhibit single foci (Fig. 2b and 2c, top; and Movie S1), while in other embryos foci were not observed (Fig. 2b and 2c, bottom; and Movie S2). The former and latter were likely to represent female and male, respectively. In the later stage embryos, the sex of embryos was determined by genomic PCR using extraembryonic membranes. In female embryos, nuclear foci were still observed in most nuclei in E10.5, but disappeared in E14.5 (Fig. 2d). To confirm the H4K20me1-mintbody foci correspond to Xi and that its concentration decreases by E14.5, we stained wild-type mouse embryos using antibodies specific for H4K20me1 and H3K27me3, which is an Xi marker (Fig. 2e and 2f). As expected, H4K20me1 was concentrated in H3K27me3-erinched foci in most cells in E7.5 (Fig. 2e), but the co-localization of H4K20me1 and H3K27me3 was less clear in E14.5 (Fig. 2f). These data support the view that H4K20me1-mintbody accurately represents the intranuclear distribution of H4K20me1.

**Fig. 2.**
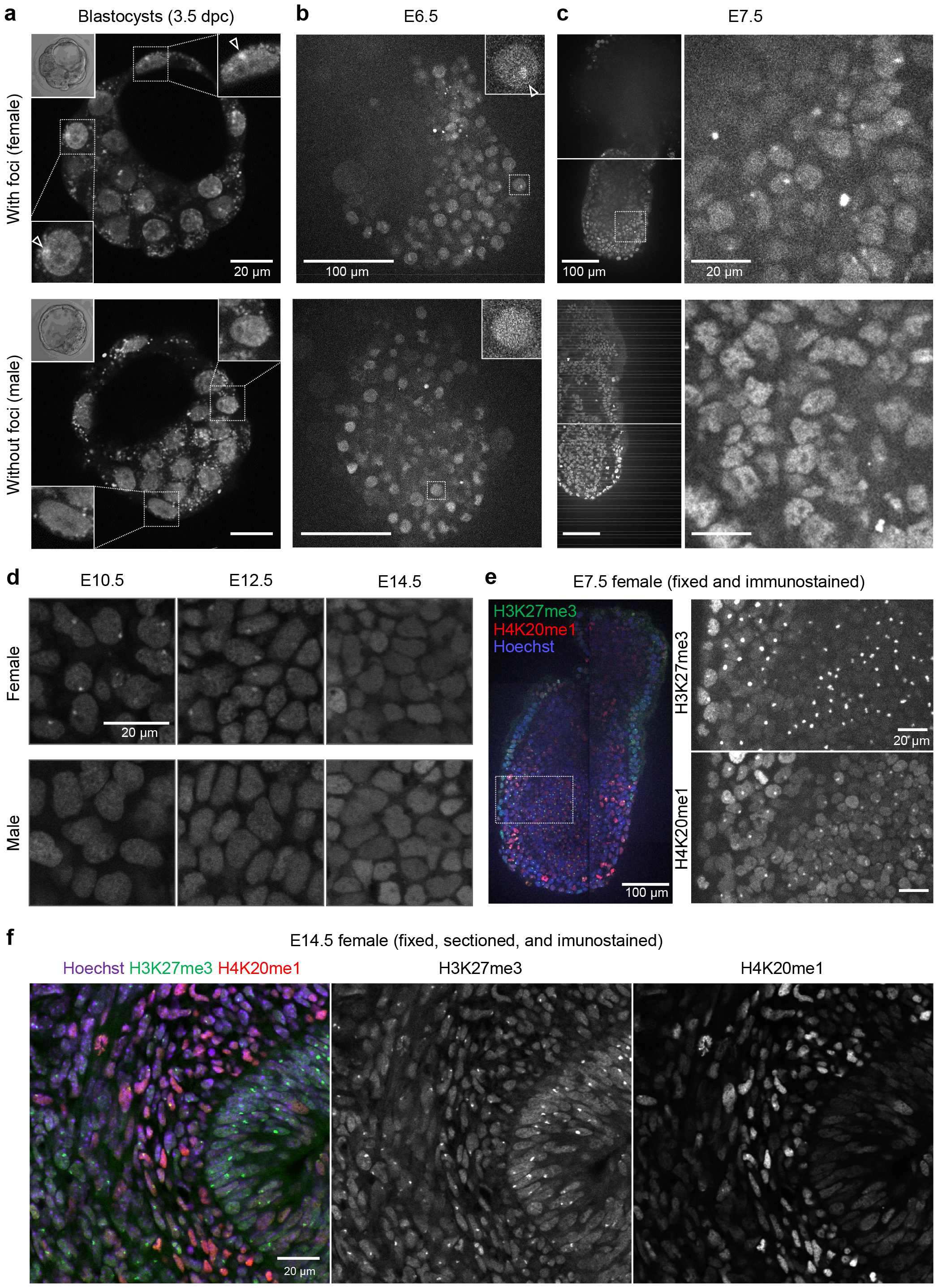
H4K20me1-mintbody in mouse embryos. (a-d) H4K20me1-mintbody in unfixed mouse embryos at the blastocyst (a), E6.5 (b), E7.5 (c), and E10.5, E12.5, and E14.5 (d) stages were visualized using a confocal microscope. (a) The 3.5 dpc blastocyst-stage preimplantation embryos. Maximum intensity projections of three confocal sections are presented alongside bright-field images of embryo (top left insets) and magnified views of selected nuclei (top right and bottom left insets). Nuclei in probable female embryos exhibit single foci (open arrowheads; top panel). (b) E6.5 embryos. Single confocal sections are shown with magnified views of selected nuclei (insets). An open arrowhead indicates a focus (top panel). (c) E7.5 embryos. Low- and high-power views of single confocal sections are shown on the left and right, respectively. See Movie S1 and S2 for the full z-stack images of probable female and male embryos, respectively. (d) E10.5, E12.5, and E14.5 embryos. The sex of each embryo was determined by genomic PCR. The number of nuclei with foci appears to gradually decrease from E10.5 to E14.5. (e and f) Embryos were fixed and stained with antibodies specific for H3K27me3 and H4K20me1. DNA was counterstained with Hoechst 33342 before acquiring confocal images. The sex of each embryo was determined by genomic PCR. (e) E7.5 female embryo. Merged and magnified single-color views are shown on the left and right, respectively. H4K20me1 foci overlap with H3K27me3 foci. (f) E14.5 female embryo section. H4K20me1 is not concentrated in H3K27me3 foci.

### H4K20me1-mintbody highlights XY bodies in pachytene cells during spermatogenesis

To demonstrate that H4K20me1-mintbody represents the dynamic changes of H4K20me1 *in vivo*, we next visualized male mouse testis because H4K20me1 distribution is reported to dynamically change during the spermatogenesis (van der Heijden et al. 2007; Wang et al. 2021). At the pachytene stage, in particular, H4K20me1 is reported to be enriched in XY body, which consists of unsynapsed X and Y chromosomes and is subject to meiotic sex chromosome inactivation. When confocal sections were acquired for the P17.5 immature mouse testis, just isolated and placed on a glass bottom dish without fixation, a variety of H4K20me1-mintbody distribution patterns were observed in different cell nuclei (Fig. 3a and Movie S3). Near the basal membrane (z = -3.0 μm), relatively large zygotene-like nuclei and smaller interphase nuclei, possibly representing spermatogonial cells or spermatocytes, were observed (1 and 2) (van der Heijden et al. 2007; Nakata et al. 2015; Ueda et al. 2017). The higher signal intensity in zygotene-like nuclei is consistent with zygotene cells being arrested at G2 phase, during with H4K20me1 level increases in somatic cells (Rice et al. 2002; Pesavento et al. 2008; Sato et al. 2016). At the deeper section (z = 33 μm), pachytene cell nuclei with single foci were observed in addition to smaller interphase nuclei (3 and 4). When a seminiferous tubule was pulled out of an immature testis, a gradual change in H4K20me1-mintbody’s subnuclear distribution during differentiation was observed along with the tubule (Fig. 3b), from dividing cells showing condensed mitotic chromosomes (1), to cells showing interphase nuclei (2), zygotene nuclei (3), and pachytene nuclei showing intense nuclear foci (4). In a 1.5-month-old mouse testis, pachytene nuclei with foci, and round and elongated spermatid nuclei were observed both in intact testis (Fig. 4a) and isolated tubules (Fig. 4b). These observations are consistent with the progression of spermatogenesis over months. To confirm whether H4K20me1-minbody foci in pachytene nuclei correspond to XY bodies, meiotic cell spreads were stained with specific antibodies directed against synaptonemal complex protein SCP3 and γ-H2AX, the phosphorylated form of H2A induced by DNA double strand breaks and a marker of XY body (Fernandez-Capetillo et al. 2003; van der Heijden et al. 2007). H4K20me1-mintbody signals were indeed enriched in γ-H2AX foci in SCP3-positive cells, although not all XY bodies exhibited H4K20me1-minbtody concentration either in P15.5 and 4.5-month-old mice (Fig. 5). Thus, the distribution of H4K20me1 in differentiating testicular cells can be accurately detected by the H4K20me1-mintbody.

**Fig. 3.**
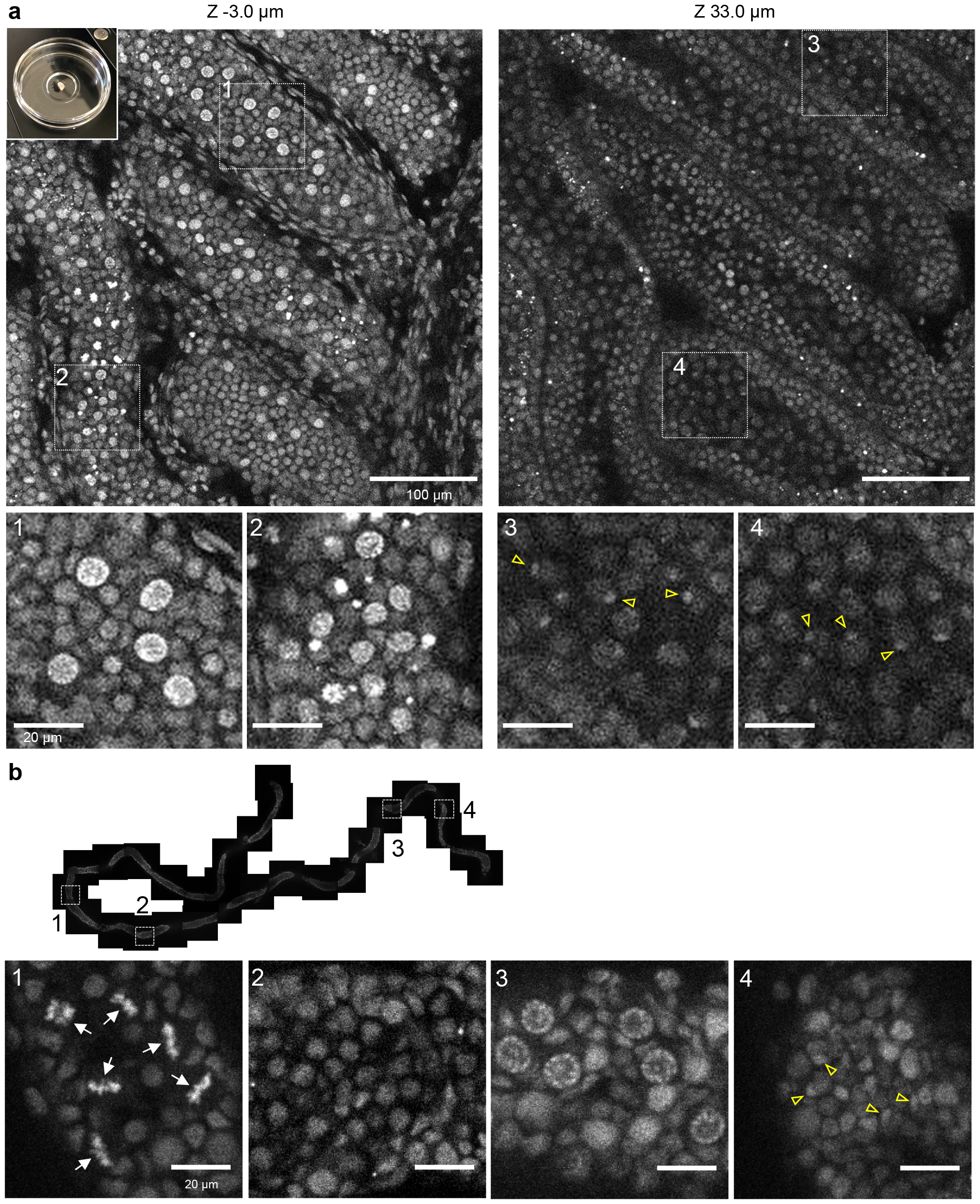
H4K20me1-mintbody in P17.5 mouse testis. (a) The testis was isolated from P17.5 male mouse expressing H4K20me1-mintbody and placed onto a 35-mm glass-bottom dish (top left inset) for confocal microscopy. Two different z-stack images with magnified views of indicated areas (1-4) are shown. See Movie S3 for the full z-stack images. (b) A seminiferous tubule was isolated from a P17.5 testis, placed onto a glass-bottom dish for confocal microscopy. Tiled views along the tubule and magnified views of indicated areas (1-4) are shown. Arrows and open arrowheads indicate mitotic condensed chromosomes and nuclear foci, respectively.

**Fig. 4.**
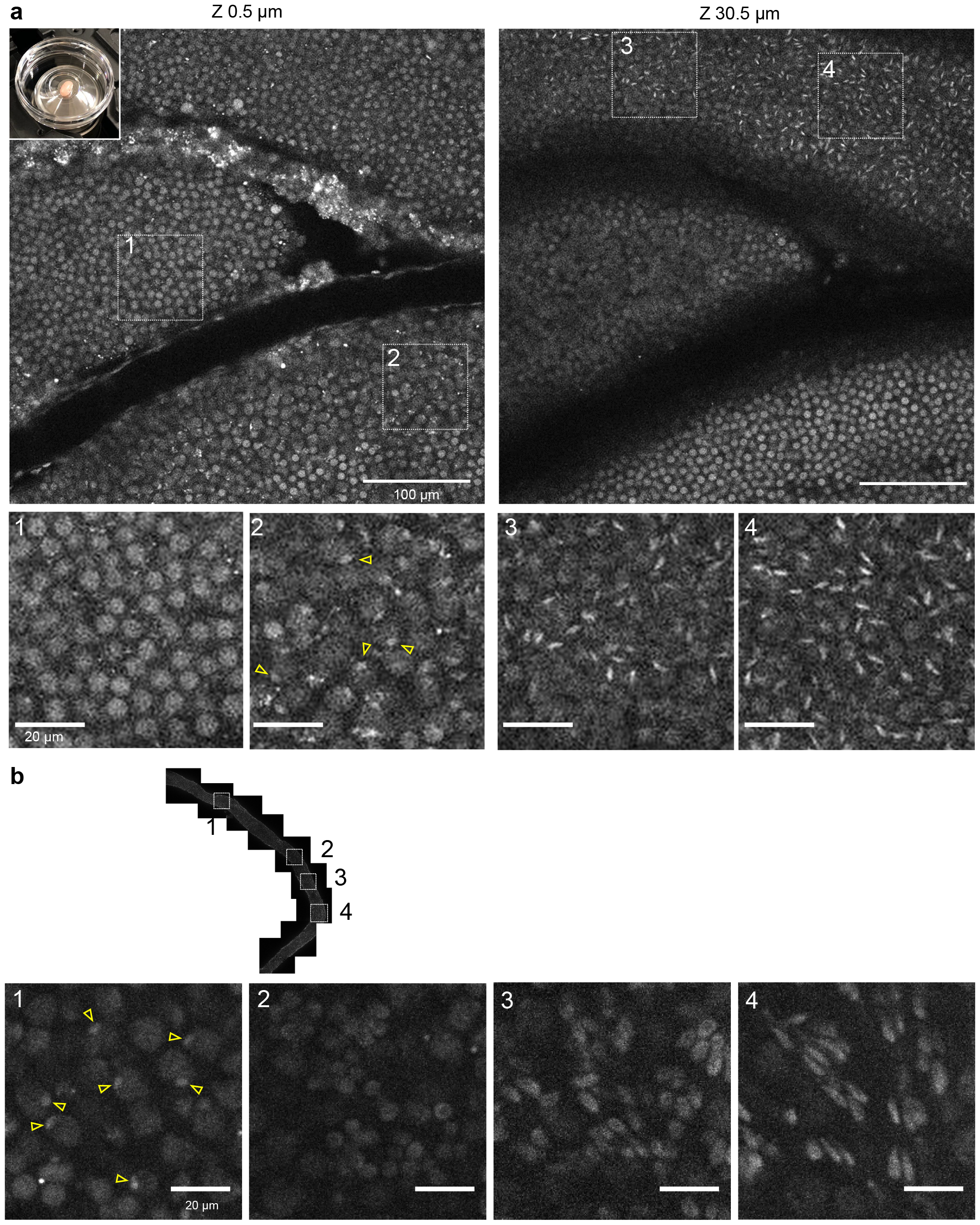
H4K20me1-mintbody in 2.5-month-old mouse testis. (a) The testis was isolated from 2.5-month-old male mouse expressing H4K20me1-mintbody and placed onto a 35-mm glass-bottom dish (top left inset) for confocal microscopy. Two different z-stack images with magnified views of indicated areas (1-4) are shown. See Movie S4 for the full z-stack images. (b) A seminiferous tubule was isolated from a 2.5-month-old testis, placed onto a glass-bottom dish for confocal microscopy. Tiled views along with tubules and magnified views of indicated areas (1-4) are shown. Open arrowheads indicate nuclear foci.

**Fig. 5.**
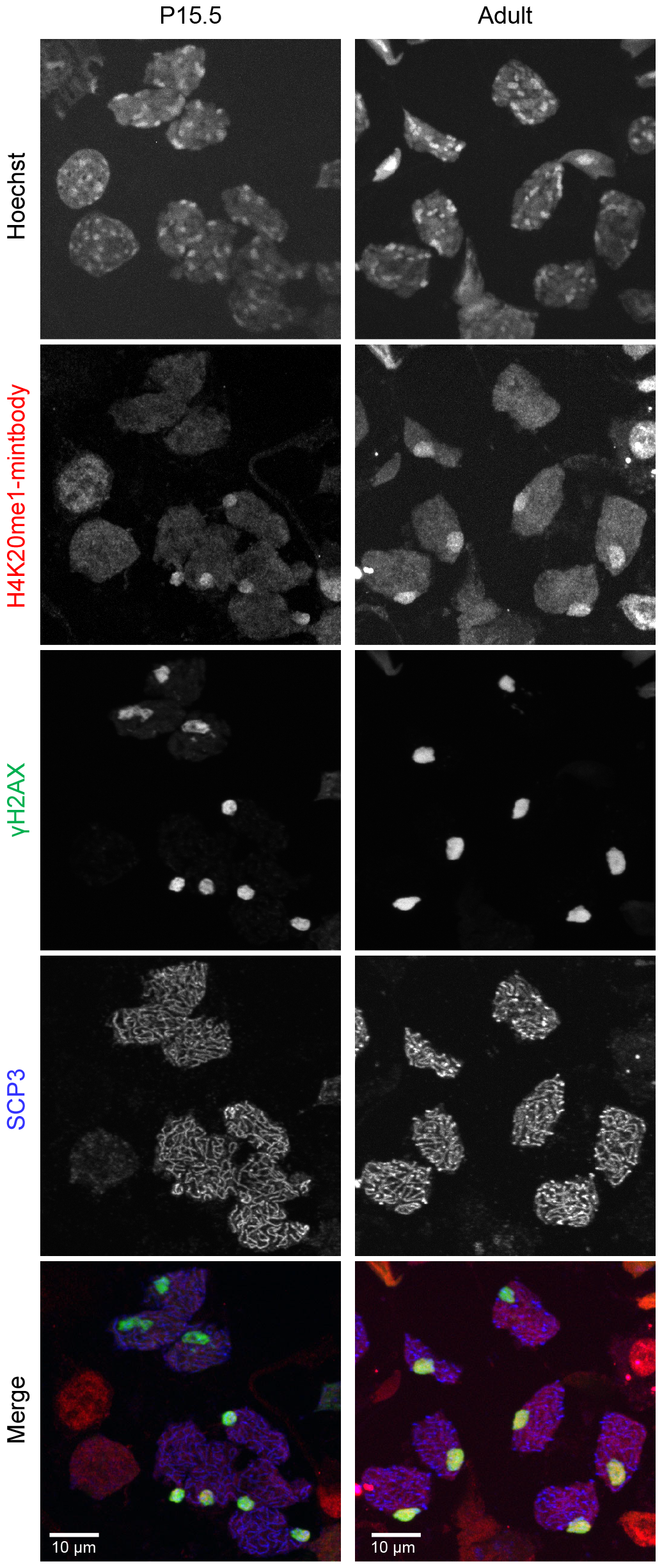
H4K20me1-mintbody enriched foci correspond to XY bodies. Cells in seminiferous tubules from P15.5 and 2.5-month-old mice expressing H4K20me1-mintbody were spread, fixed, and stained with antibodies specific for γH2AX and SCP3. Max intensity projection images of confocal sections are shown.

## Conclusions

We demonstrate here that H4K20me1-mintbody allows us to visualize the localization of H4K20me1 in mouse tissues without the need for fixation. While any protein can, in principle, be visualized in mice by tagging with a fluorescent protein, visualizing posttranslational modifications *in vivo* has been challenging. Live-cell probes that detect specific modifications, such as mintbodies, have addressed this issue, yet only non-mammal organisms like Drosophila and C. elegans expressing mintbodies have been developed. This is the first report to show that mammals expressing the mintbody throughout the entire body developed normally and were fertile. Although any ectopic probe that binds to specific modification could interfere with the binding of endogenous proteins, this suggest that the epigenetic readout system during development and differentiation is tolerant to a certain level of expression of a low-affinity binding probe. Mice expressing H4K20me1-mintbody will be useful for future work on the dynamics of this modification in mice and also ex vivo histochemical analysis.

## Acknowledgements

The mouse line was generated with the support of the Institute for Microbial Diseases, Osaka University, and we thank Mitsuko Mori and Masahito Ikawa for their assistance.

## Author contributions

Conceptualization, Y.S., H.K.; investigation, Y.S., M.T., M.U., J.U.; methodology, Y.S., J.U., K.Y.; writing – original draft, H.K., writing – review and editing, Y.S., M.T., M.U., J.U., K.Y.; funding acquisition, K.Y., H.K.; supervision, K.Y., H.K.

## Declarations

## Supplementary Movies

**Movie S1, H4K20me1-mintbody in E7.5 female embryo**

Confocal sections of an E7.5 female embryo expressing H4K20me1-mintbody across 60 μm at 1 μm intervals.

**Movie S2, H4K20me1-mintbody in E7.5 male embryo**

Confocal sections of an E7.5 male embryo expressing H4K20me1-mintbody across 60 μm at 1 μm intervals.

**Movie S3, H4K20me1-mintbody in P17.5 mouse testis**

Confocal sections of an P17.5 mouse testis expressing H4K20me1-mintbody across 72.5 μm at 0.5 μm intervals.

**Movie S3, H4K20me1-mintbody in 2.5-month-old mouse testis**

Confocal sections of an adult mouse testis expressing H4K20me1-mintbody across 84.0 μm at 0.5 μm intervals.

